# Fatigue Metabolites of Large Yellow Croaker and Construction of Swimming Model

**DOI:** 10.1101/2021.12.01.470794

**Authors:** Ruoyu Chai, Heng Yin, Runming Huo, Xiaomei Shui, Hanying Wang, Ling Huang, Ping Wang

**Affiliations:** National Engineering Research Center for Facilitated Marine Aquaculture, Zhejiang Ocean University, Zhoushan 316022, China

**Keywords:** aquaculture, large yellow croaker, swimming ability, flow rate preference, glycogen, lactic acid

## Abstract

A trend in large yellow croaker (*Larimichthys crocea*) aquaculture is to establish new production sites that are suitable for extreme weather conditions. However, continuous and strong currents can harm fish welfare. To determine the location of the net cage, it is necessary to assess the swimming ability of large yellow croaker. Currently, our research on large yellow croakers is focusing on behavior analysis. This article investigates the effect of swimming large yellow croakers on metabolites in the body by examining the preferred speed of the group and the endurance swimming ability of single-tailed fish. We evaluated the factors that influence the large yellow croaker’s swimming fatigue by quantifying the content of metabolites and constructed the endurance swimming model using those results. Various results showed large yellow croaker populations tend to grow in low-velocity environments, and this matches their traditional habitat. The samples were taken at different swimming times at a flow rate of 0.35 m/s. According to the results of the metabolite content determination, blood glucose levels is closely related to swimming ability in large yellow croakers. The content of liver glycogen, which regulates blood glucose concentration, decreases in a certain linear relationship. The endurance swimming model of large yellow croaker was constructed according to the changes of liver glycogen content. The goals of this article are to provide a deeper understanding of the physiological characteristics of large yellow croaker swimming, and to provide a reference for choosing fishing and cage sites for large yellow croaker.

## 1. Introduction

The large yellow croaker was once an endemic commercial marine fish in East Asia[1]. The wild population of this fish has been severely depleted since 1950 due to heavy fishing[2]. China has taken active protection measures to protect large yellow croakers since 1980[3], including breeding and releasing, fishing bans, and habitat maintenance[4]. The large yellow croaker is mostly found around the coasts of East Asia, especially from the southern Shandong Peninsula in the north to the western Taiwan coast in the east[2, 5–7]. In 2019, the aquaculture production reached 225,549 tons, which is the most productive fish species in China’s marine aquaculture[8]. There is evidence that wild large yellow croakers move seasonally[2, 9]. In the spring and autumn, large yellow croakers lay their eggs in shallow coastal waters (<30 m). The water temperature drops in winter, and the fish gather in deeper waters (50-80m offshore)[10]. The migration of large yellow croakers is also affected by temperature, as is that of sockeye salmon[11]. Large yellow croaker usually needs to move tens to hundreds of kilometers. The powerful currents challenge their swimming ability.

Currently, the large yellow croaker aquaculture industry is trying to build deep-water cages in the ocean that can survive extreme weather (such as strong winds and huge waves)[12, 13]. Technology advancements and novel management strategies are required for this process. Furthermore, it is vital to determine whether the fish can thrive in a harsher environment. Continuous currents are considered the greatest threat to fish welfare[14]. Large yellow croakers farmed in cages in low currents will swim around the cage or hit the purse seine at the speed of the fish[15]. However, large yellow croaker will stop cruising when its current speed exceeds its current speed[16, 17]. The literature shows that voluntary actions are restricted, and mandatory increases in flow rates may undermine fish welfare. Additionally, it could limit the fish’s growth because it requires a lot of energy to carry out continuous swimming[18, 19]. Therefore, it is crucial to understand the swimming ability of farmed fish in order to scientifically select suitable sea areas for aquaculture.

The ability to swim is vital for the survival of fish. By swimming, fish accomplish several activities necessary to survival, including migration, evading predators, and escaping predation[20, 21]. Flow rates that are appropriate ensure good water quality, prevent ammonia nitrogen build-up in the water, and the spread of fish diseases as well as supporting fish growth and metabolism[22]. The research on fish swimming ability is relatively extensive at present. According to the different swimming speeds of fish, it can be divided into Sprint swimming speed, Critical swimming speed[23], and Maximum sustainable swimming speed[24]. While some models of fish swimming speed and fatigue have been partially accepted, they have not been fully explained based on physiology[25]. The current research on the swimming behavior of large yellow croakers includes endurance swimming time and critical swimming speed[26].

The currents in cage aquaculture can be so intense that they persist for hours or days due to extreme weather, and burst swimming can be so short that it lasts only minutes. Therefore, it is essential to assess large yellow croakers’ continual swimming ability, which is known as “high-intensity swimming.” This system is entirely powered by red aerobic slow muscle fibers, while white aerobic fast fibers remain inactive[27, 28]. Contrary to burst swimming speeds, continuous swimming should not cause fatigue since it avoids lactic acid accumulation and maintains homeostasis. But lactic acid content is not the only factor that contributes to fish swimming fatigue. For example, the physiological stress caused by exhaustive exercise can be severe enough to cause animal death a few hours later[29].

For practical reasons, continuous swimming is usually defined as being able to maintain a speed of 200 minutes[30]. The literature shows that the distance that fish can swim in areas with high-speed currents is an important factor that limits the distribution, protection, and cage culture of river fish and amphibians[31]. Rebecca Fisher investigated the critical swimming speed of the juvenile coral fish (*Alcyoneum piscis*) and found that its swimming speed is affected by factors such as habitat and morphology[32]. The critical swimming speed of the black sea bream (*Sparus macrocephlus*) with a size of 215.5 g is about 0.65 m s^-1^, while the American redfish (*Sciaenops ocellatus*) with a size of about 186.9 g can still maintain 200 min at a flow rate of 0.80 m s^-1^. In this comparison, it can be seen that the American redfish is very good at swimming. The difference in swimming ability among various fish species is primarily due to body surface characteristics and metabolism. Zhu Yanping’s research on *Pelteobagrus vachelli* showed a link between fish swimming ability and metabolites in the body, showing that muscle and blood lactic acid levels increased over time while liver, muscle, and blood glycogen levels declined[33]. In contrast, Chao Shuai’s study of metabolites in American redfish demonstrated that blood glucose and lactic acid levels increased without a significant change in muscle glycogen levels[34]. Research shows that due to the combined effects of metabolic and respiratory acidosis, the blood pH of all fishes is significantly lowered following exhaustive exercise. However, in continuous aerobic exercise, the blood acid-base state has little change compared with the initial state[35]. The comparison reveals that fish species differ in their metabolisms. The results of actual experiments on the large yellow croaker must be discussed further based on the results.

In this study, continuous experiments were conducted on three items related to large yellow croaker swimming speed preference, endurance swimming ability, and the relationship between swimming ability and metabolites in the body. On the basis of the experimental data, the key factors that cause fatigue are identified by excluding the influence of environmental factors like the ammonia nitrogen content of the water and the temperature[36, 37]. The results may represent a new reference for determining sea area selection for breeding cages for large yellow croakers and a new method for recycling water for factory breeding, improving the efficiency of breeding enterprises.

## 2. Materials and Methods

### Ethics Statement

This study was conducted in strict accordance with the laws governing animal experimentation in China. The protocol was approved by the China Zhejiang Ocean University.

### 2.1 Experimental Materials

The experimental fish were purchased from the breeding station in Zhoushan(site A and site B, figure 1), Zhejiang Province, China (30°02′ N, 122°13′ E). In the laboratory, 100 large yellow croakers with an initial weight of (28.4±0.9) g were trained for two weeks in a water tank with automatic oxygen pumping. At the time of training, the water temperature was constant at (20±1) °C, the dissolved oxygen in the water was greater than 6.5 mg L^-1^, the photoperiod was 12L:12D, and commercial pellets were fed at 8:00 and 19:00 daily. Feeding amounts comprise 3% of the fish’s weight in the tank. Furthermore, after two hours of bait feeding, we removed the residual baits and feces, and exchanged one-third of the water each day. The experimental water body and the temporary breeding water body remained the same during the temporary breeding period. The experimental fish were anesthetized with clove oil at a concentration of 60mg/L and sacrificed. All efforts were made to minimize suffering.

**Fig.1.**
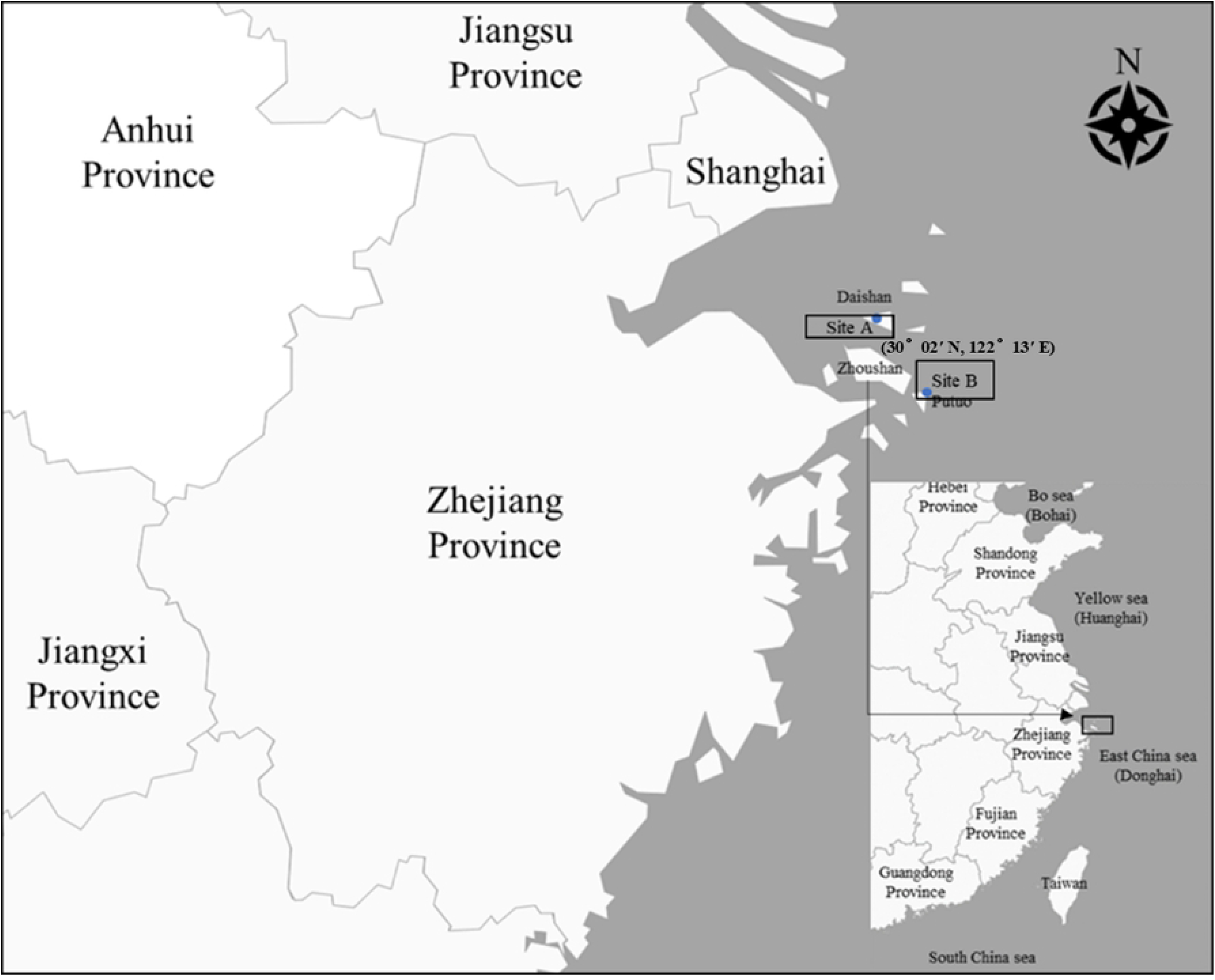
Fishing site location.

### 2.2 Laboratory equipment

In this experiment, the air pump provides sufficient oxygen during the experiment, and the dual-body water tank provides the multi-functional environment required. An electric propeller and mesh rectifier are used to create a stable and uniform circulating water flow to test the ability of large yellow croakers to swim. The Vectrino point-type flow meter was placed in the center of the water channel, and the probe was 15 cm from the bottom of the water tank in order to measure the water flow speed in real-time. The camera (Nikon, 25 frames s^-1^) is set up on the side of the observation area, the lens is 80 cm away from the side glass, and the whole process of the large yellow croaker swimming and touching the net is documented. This tank has a total length of 17.13 meters, divided into two tanks of 10 meters each. It is used to test the endurance swimming ability of large yellow croaker. One of them, tank 2, does not change in cross-sectional area, and the water flow speed is the same everywhere. Research has revealed that the choice of experimental equipment has a greater impact on the swimming ability of fish. Therefore, a metal cage is set up in the water tank to simulate a net structure to achieve a conventional cage culture environment. Over the course of tank 1, the cross-sectional area of tank 1 gradually decreases. A is 80×80 cm, and B is 20×80 cm. This flows at a velocity of *U_B_*=4*U_A_*. U_A_ and U_B_ are the velocities of the water moving through A and B, respectively, to generate a flow gradient in a tank. Clusters of large yellow croakers select the flow velocity and examine their swimming preferences, as shown in Figure 2.

**Fig.2.**
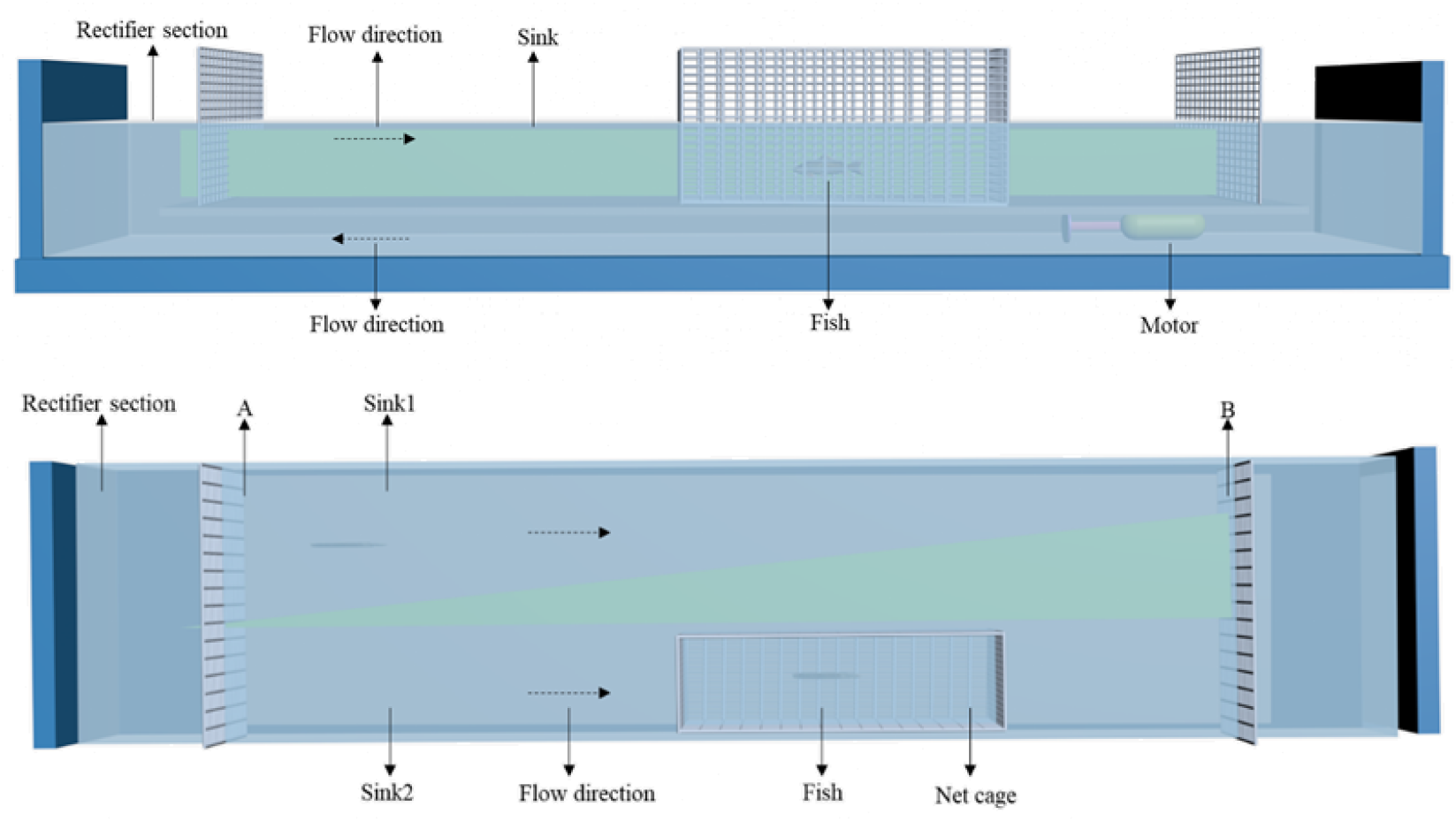
Diagram of an experimental water tank. The rotating impeller maintains the water flow.

### 2.3 The swimming speed preference of large yellow croaker

Take 64 fish and place them in tank 1, and set 5 sets of flow rates(1: 0.024~0.096 m s^-1^;2: 0.035~0.140 m s^-1^;3: 0.050~0.200 m s^-1^;4: 0.078~0.312 m s^-1^;5: 0.095~0.380 m s^-1^). From point A, a low-velocity area is at a distance of 0 to 3 m, and a high-velocity area is at a distance of 7 to 10 m. First, the fish were placed in a static water flow tank for more than 24 hours to adjust, and then they were adjusted to reach 0.024 m/s within 1 minute. Monitor and record the distribution of fish in the aquarium with an external camera that lasts for 200 minutes, and records the distribution of fish in different areas. After turning off the engine for 60 minutes, adjust the flow velocity to 0.035 m s^-1^, 0.05 m s^-1^, 0.078 m s^-1^, and 0.095 m s^-1^ for a total of 5 experimental groups. Repeat the experiment as above. Record the distribution of large yellow croakers in the tank at the flow rate of each group for 200 minutes. At the end of each group of experiments, the statistics do not include the fish caught in the tank and swimming back and forth.

### 2.4 The endurance swimming ability of large yellow croaker

In the water tank, the fish behave in ways that interfere with the experiment, such as sticking to the net and swimming against the wall. In the experimental water tank, a metal cage is set up to conduct swimming experiments. The preliminary experiment determined the water flow velocity of six experiments, which were 0.15 m s^-1^, 0.25 m s^-1^, 0.35 m s^-1^, 0.4 m s^-1^, 0.45 m s^-1^, 0.5 m s^-1^. The speed of the flow of water is the speed of swimming of the experimental fish against the current.

The large yellow croaker stopped feeding one day before the endurance swimming trial, and selected subjects with similar characteristics for the experiment. The swimming experiment was repeated 6 times, each time with a different fish. This study does not repeat experiments since it is difficult to guarantee the same level of recovery for large yellow croaker after fatigue. The experiment proceeds as follows: Set up the camera, then place the trial fish in tank A for at least 60 minutes to adapt. After one minute, adjust the flow rate to the preset value and record the cumulative swimming time of each experimental fish at this rate until it becomes fatigued. The criterion for judging swimming fatigue is when the experimental fish stop swimming and touch the net for at least 20 seconds. Weigh the swimming body using an electronic balance (accurate to 0.01g) and measure its weight and length while recording the continuous swimming time. In this experiment, we measured the swimming time and the swimming speed and calculated the swimming ability index (SAI)[38]. The SAI refers to the area of the graph enclosed by the curve of swimming time and swimming speed and the coordinate axis. In addition to swimming speed, it also measures swimming time, so it is a good indicator. It can also be used to compare swimming ability between species or at different growth stages of the same species. The calculation formula is: 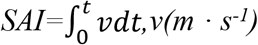 is the swimming speed, t(s) is the swimming time.

### 2.5 The relationship between the swimming ability of large yellow croakers and their metabolites

Based on the results of the endurance experiment, the water tank’s flow velocity and level are set to 0.35 m s^-1^ and 0.40 m, respectively. Large yellow croakers with similar specifications and fasting for 12 hours were randomly selected and placed into the water tank. At each of six time points—0, 30, 60, 90, 120, and 150 minutes—six fish were randomly selected for tail-by-tail testing. The experiment was started after the fish had been adapted for 60 minutes at a flow rate of 5 cm s^-1^. Within one minute, adjust the flow rate of the water tank to 0.35 m s^-1^.

Inject 200μL of blood into a 1 mL centrifuge tube using a pre-cooled sterile syringe, and mix it with the collected sample. Centrifuge at 3000 r min^-1^ at 4°C for 10 minutes in a centrifuge. Transfer the supernatant into a centrifuge tube and freeze it at −20°C. During the same procedure, liver and muscle were removed from the fish and stored at −80 °C. Each experiment guarantees at least five effective fish. It was determined that liver glycogen, muscle glycogen, blood lactic acid, muscle lactic acid, and fish glucagon were all metabolites. All reagents have been provided by Nanjing Jiancheng Institute of Biological Engineering.

### 2.6 Statistics

The experimental data are routinely calculated using Excel 2019, and then SPSS 26.0 is used for statistical analysis. An independent sample t-test is used to analyze the physiological index differences between the experimental fish prior to and after exercise fatigue in the endurance swimming experiment. We use one-way analysis of variance along with multiple Duncan comparisons to analyze different swimming speeds. In this research, we calculated the relationship between the swimming speed and sustainable swimming time using curve regression, with a significance level of P<0.05. SPSS 26.0 statistical software is used for all data processing. In the results, the mean is multiplied by the standard error.

## 3. Results

The weight, body length, and body height of each treatment group are shown in Table 1. All fish came from the same holding tanks and were tested Table 1. Size parameters for the large yellow croaker swimming trials, showing mean±SE weight, body length, height and number of fish measured (n) in each treatment group. Different superscript letters indicate a significant difference (p < 0.05).

**Table.1.** The morphological parameters of experimental fishes under different swimming speed

$n=5; \bar{x} \pm SD$
| Measure data | Swimming Speed (cm s<sup>-1</sup>) |  |  |  |  |  |
| --- | --- | --- | --- | --- | --- | --- |
|  | 15 | 25 | 35 | 40 | 45 | 50 |
| Weight | 27.93±2.64<sup>a</sup> | 27.8±3.76<sup>a</sup> | 28.5±1.25<sup>b</sup> | 27.75±2.19<sup>a</sup> | 28.12±1.29<sup>ab</sup> | 27.96±2.51<sup>ab</sup> |
| Length | 13.61±0.19<sup>a</sup> | 13.58±0.13<sup>a</sup> | 13.63±0.23<sup>a</sup> | 13.3±0.01<sup>a</sup> | 13.63±0.07<sup>a</sup> | 13.77±0.02<sup>a</sup> |
| Hight | 3.69±0.02<sup>a</sup> | 3.66±0.03<sup>a</sup> | 3.68±0.04<sup>a</sup> | 3.75±0.01<sup>b</sup> | 3.67±0.01<sup>a</sup> | 3.73±0.02<sup>ab</sup> |

Note: Values in each row without a common superscript are significantly different ( $P < 0.05$ )

### 3.1 Swimming speed preference experiment results

We define the two behaviors of the fish in the flow rate selector and the human’s selection of the respiratory rate as preferences. According to Schizothorax oconnori exhibits upstream-swimming and searching in high-speed flow conditions [39]. The results of Wang indicate that the critical swimming speed of large yellow croaker is 0.5 ± 0.02 m·s-1 at 19.0~ 21.0°C [40]. The flow velocity range of the flume designed in this test is 0.024-0.380 m·s-1, and the maximum flow velocity is less than 0.5 m·s-1. The water tank did not set up obstacles to the swimming back and forth of the test fish. It should also be noted that this study provides a variety of flow rate environments for voluntary swimming, which is significantly different from the setting of the swimming ability experiment. Therefore, the residence time of large yellow croaker in the tank can be attributed to the preference for flow rate.

Figure 3 indicates that the large yellow croaker cluster are relatively stable at different flow rates. The distribution of fish in areas below 0.1 m s^-1^ did not differ significantly (P<0.05). It shows that large yellow croaker populations like to survive in low-velocity environments, especially in clusters below 0.1 m s^-1^.

**Fig.3.**
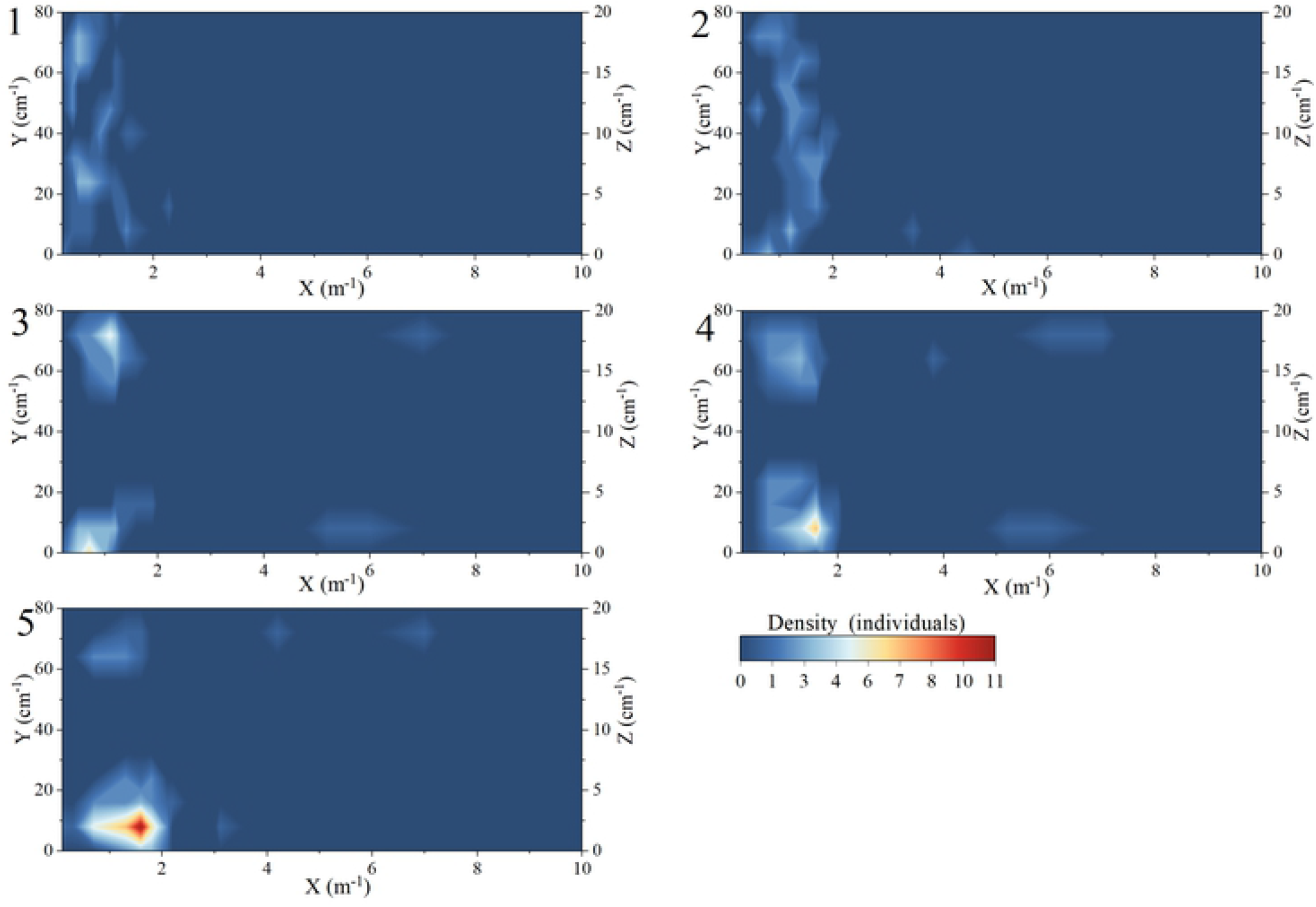
Changes in fish cohesion during a swimming preference test. The X-, Y and Z - axes indicate horizontal position along the lengthwise and widthwise axes of the swim chamber respectively. The left axis indicates the front of the swim chamber.

### 3.2 Endurance swimming ability experiment result

The endurance swimming time of large yellow croaker gradually decreases with the increase of swimming speed. To study the relationship between the performance level of large yellow croakers with the flow rate, we used the average value of all large yellow croakers’ endurance swimming time at each flow rate level as the performance level with the flow rate. Figure 4 shows the measurement result as a hollow triangle. As the flow velocity increases, the sustainable swimming time decreases, but it does not change linearly. Instead, it decays more rapidly in the low-flow area and slower in the high-flow area.

**Fig.4.**
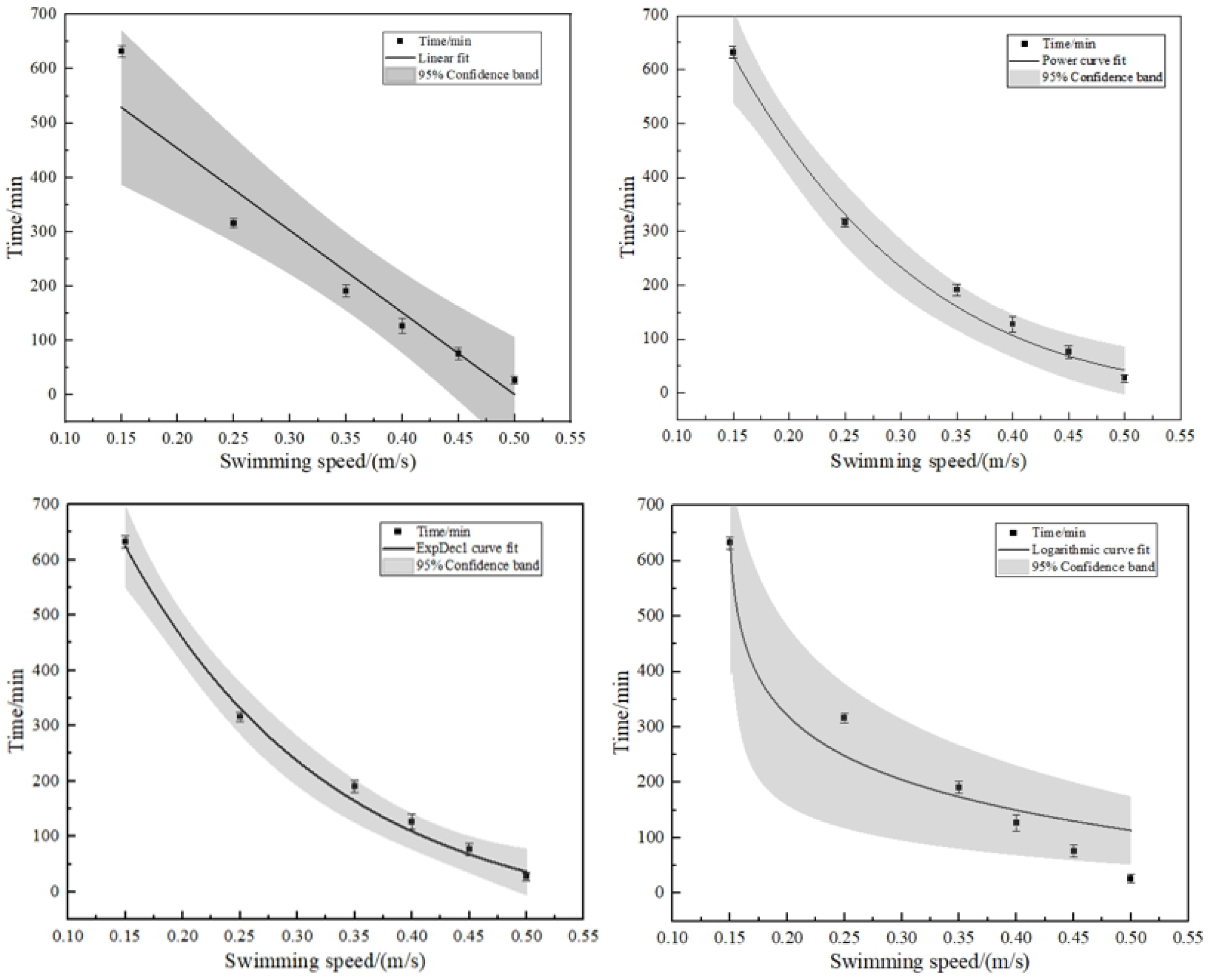
Relationship between average endurance swimming time and flow velocity. Vertical bars represent standard deviations. Grey part represents 95% confidence band.

Based on the above swimming characteristics and the curve characteristics of each function, four mathematical models of linear function, exponential function, power function, and logarithmic function are used to fit the relationship curve between endurance time and flow velocity. In Figure 4, the solid line represents the result. In Table 2, we find the fitting equation and the determined coefficient R^2^. With the value of R^2^, it can be determined the powerful function and exponential function represent the closest regression relations between endurance swimming time and swimming speed (R^2^>0.99). However, the R^2^ indicator only indicates how well the model fits the data, and cannot tell whether the constructed model is accurate. Model judgment should be measured by Akaike’s information criterion (AIC) for each of the four fitting curves[41]. The AIC is a model that assesses how two sets of data fit some degree, it’s a mix of R^2^ and adjusted R^2^. In essence, the lower the AIC, the better the model will fit the data and how to avoid overfitting. Among these models, the linear model has the best AIC (38.27). The equation is:

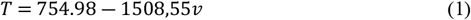

**Tab.2.** Functional relationship between the average sustainable swimming time of large yellow croaker and flow velocity

| Regression model | Fitting equation | $R^2$ | AIC |
| --- | --- | --- | --- |
| Linear fit | $T = 754.98 - 1508.55v$ | 0.930 | 38.27 |
| Power curve fit | $T = 1458.63\vert (v - 0.99)\vert ^{5.05}$ | 0.992 | 54.97 |
| ExpDec1 curve fit | $T = 1576.78e^{\frac{-v}{0.18}} - 62.40$ | 0.994 | 53.24 |
| Logarithmic curve fit | $T = -109.35 \ln(v - 0.15)$ | 0.890 | 40.49 |

Note: $T$ , swimming time; $v$ , swimming speed

According to this formula, the swimming ability index (SAI) of large yellow croakers at water temperatures of (20±1.0) °C is 188.92.

### 3.3 The changes of physiological indexes of large yellow croaker

In Table 3, we show the index changes of energy metabolites associated with large yellow croaker fatigue at different flow rates. Large yellow croaker’s glycogen and lactic acid levels changed greatly after fatigue compared with their initial levels. The contents of liver glycogen and muscle glycogen decreased significantly (P<0.05) while glucagon, muscle lactic acid, and blood lactic acid increased significantly (P<0.05). In all five groups of metabolites, the changes were significant between adjacent periods in each group (P<0.05). Figure 5 shows the change in metabolite levels in the body over time.

**Tab.3.** The content of metabolites in three tissues of large yellow croaker at different swimming speeds

| metabolite | Swimming speed (cm s<sup>-1</sup>) |  |  |  |  |  |
| --- | --- | --- | --- | --- | --- | --- |
|  | 0 | 30 | 60 | 90 | 120 | 150 |
| hepatic glycogen (mg g<sup>-1</sup>) | 17.84±0.47<sup>a</sup> | 15.76±0.51<sup>a</sup> | 10.1±0.8<sup>b</sup> | 7.22±0.35<sup>c</sup> | 4.4±0.61<sup>d</sup> | 1.74±0.19<sup>e</sup> |
| muscle glycogen (mg g<sup>-1</sup>) | 7.8±0.44<sup>a</sup> | 7.21±0.75<sup>ab</sup> | 3.84±0.36<sup>bc</sup> | 2.27±0.33<sup>cd</sup> | 1.16±0.11<sup>d</sup> | 0.38±0.11<sup>e</sup> |
| fish glucagon (ng/l<sup>-1</sup>) | 109.16±2.63<sup>a</sup> | 111.16±2.91<sup>a</sup> | 149.17±5.68<sup>b</sup> | 173.8±9.28<sup>bc</sup> | 190.76±7.07<sup>cd</sup> | 250.38±17.17<sup>cd</sup> |
| muscle lactic (mmol gprot<sup>-1</sup>) | 0.70±0.09<sup>a</sup> | 0.72±0.09<sup>ab</sup> | 1.13±0.03<sup>bc</sup> | 1.25±0.02<sup>cd</sup> | 1.42±0.04<sup>de</sup> | 1.67±0.05<sup>e</sup> |
| blood lactic (mmol l<sup>-1</sup>) | 1.31±0.08<sup>a</sup> | 1.56±0.14<sup>a</sup> | 3.00±0.13<sup>b</sup> | 4.01±0.17<sup>c</sup> | 4.63±0.06<sup>c</sup> | 5.91±0.26<sup>d</sup> |

Note: Values in each row without a common superscript are significantly different ( $P<0.05$ )

**Fig.5.**
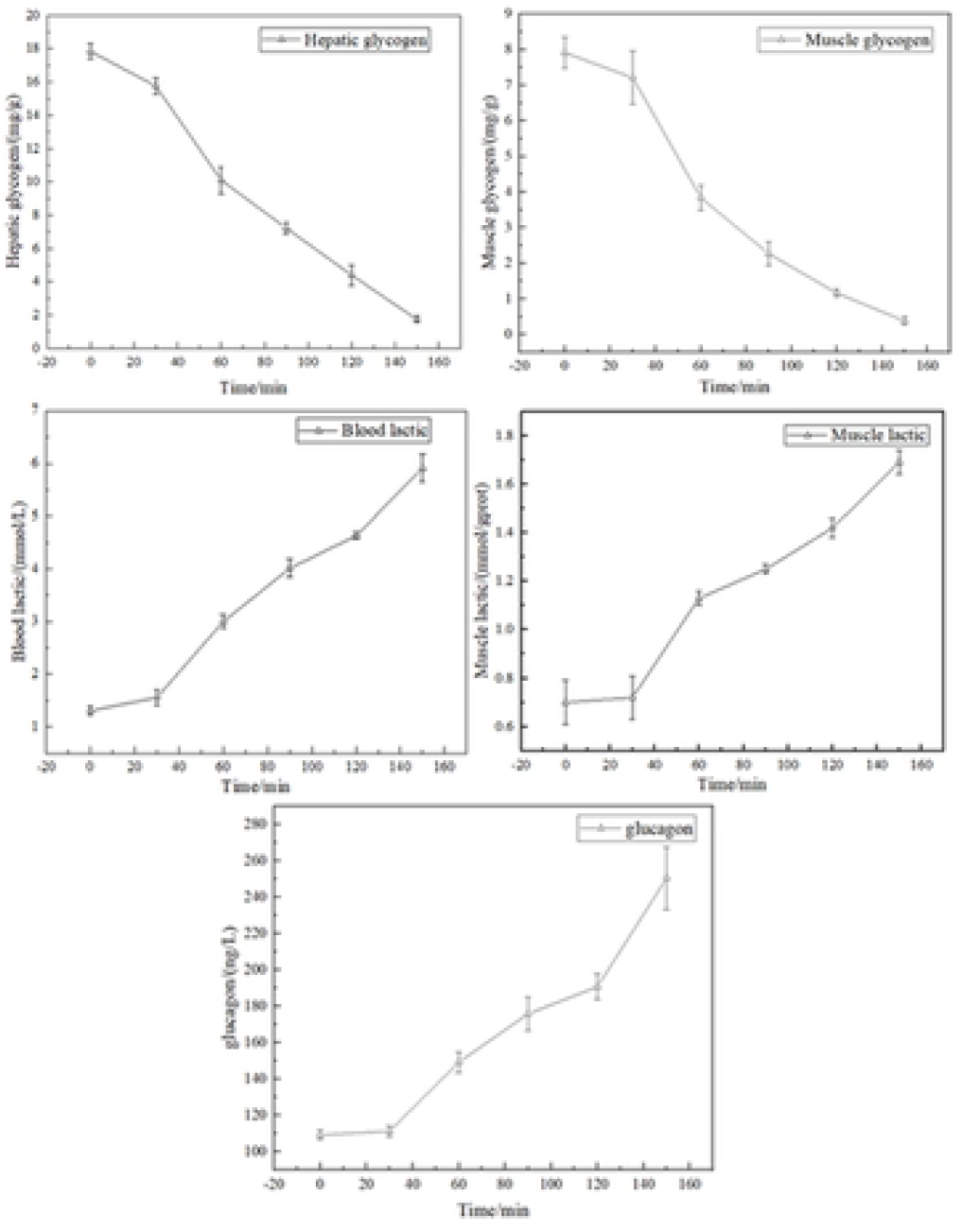
Five different metabolites change with time in the large yellow croaker. The triangle is the average metabolite concentration at this time. Vertical bars represent standard deviations.

As a result of principal component analysis (PCA). Figure 6 indicates that under the condition of the first principal component, the content of lactic acid and glycogen in large yellow croaker did not change significantly after swimming fatigue (P>0.05), while the content of glucagon was significantly different from the changes in the four groups (P<0.05). According to the data, the main factor affecting fatigue in large yellow croakers can be attributed to changes in biological blood sugar content, glucagon. As a result of the second principal component, it is obvious that the contents of the two groups of lactic acid and the two groups of glycogen respectively tend to change at the same rate, but there is a significant difference in the rate of change between lactic acid and glycogen. According to the analysis of the 6 sampling time points for this experiment, the differences in the changes in the content of the five groups of metabolites can be divided into two groups. These are the sampling time of 0, 30 minutes and the sampling time of 60, 90, 120, and 150 minutes, respectively, and the difference between 30 minutes and 60 minutes is the largest.

**Fig.6.**
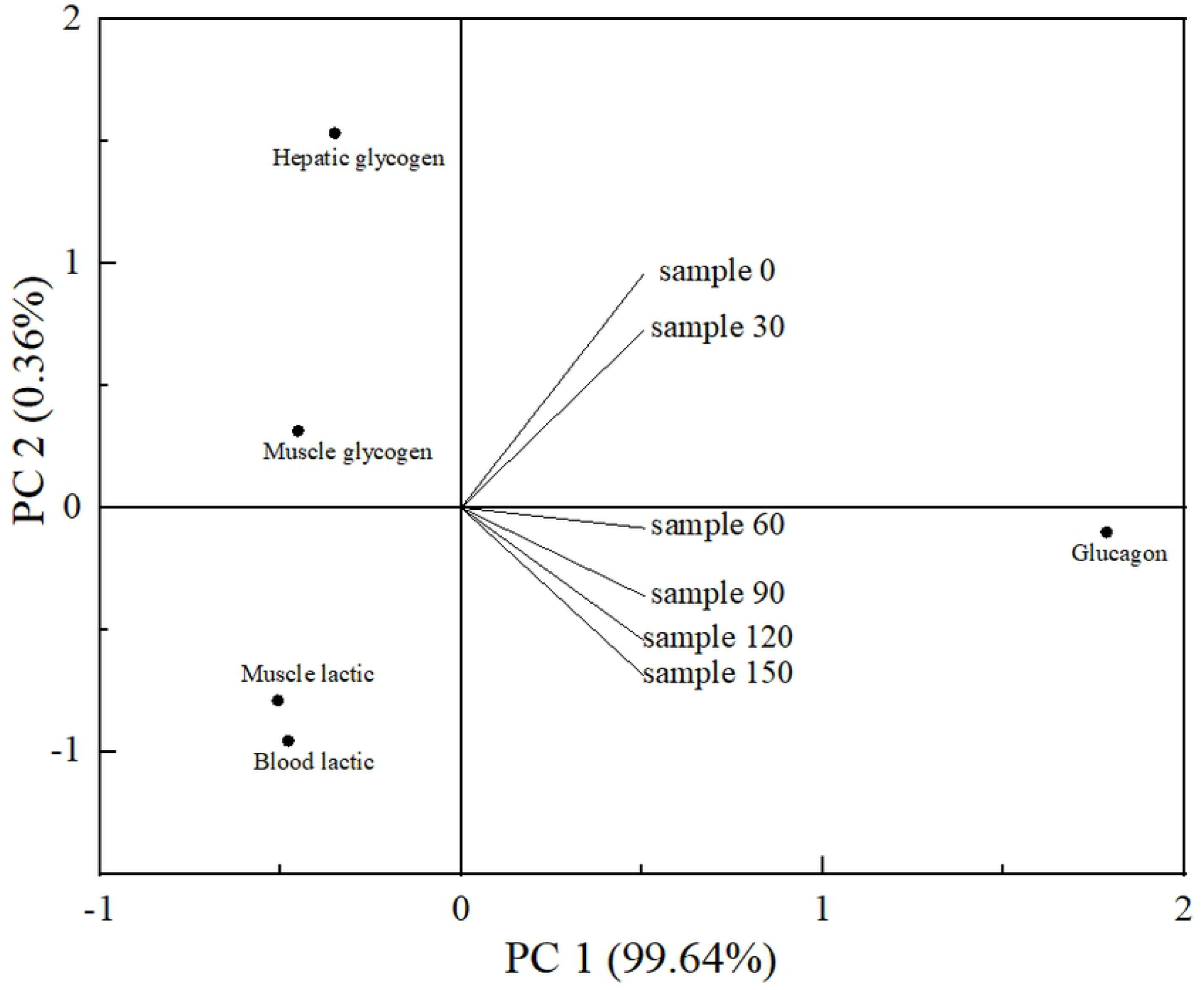
Results of principal components analysis (PCA) biplots showing relationships between five metabolites and six swimming time points. Red dots indicate the relationship between changes in the content of related metabolites.

### 3.4 The endurance swimming model of large yellow croaker

In large yellow croakers, blood sugar content regulates swimming fatigue. In this experiment, blood glucose is a key factor. An energy allocation model of large yellow croaker swimming was constructed using the variation of blood sugar content during swimming. The large yellow croaker uses some of the energy it produces from energy consumption to fight swimming resistance and some of the energy is used to meet its own needs.

The polysaccharide glycogen is known for storing energy and being readily oxidized. The liver and muscles are the main organs that store glycogen in the body. The liver glycogen accounts for about 7-10% of the wet weight, while the muscle makes up about 1-2%. The muscle’s total weight is much higher than that of the liver, so its glycogen storage is greater[42]. Muscle glycogen mainly provides energy for exercise, liver glycogen mainly maintains blood sugar levels in the body, and plays a crucial role in maintaining body health[43, 44]. Construct a fitting curve model for liver glycogen content:

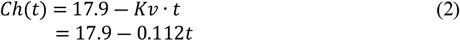

Ch(t) is the liver glycogen concentration in large yellow croaker at time t (mg g^-1^); *Kv* is the decay rate and t is time.

In the experiment, large yellow croaker was used that averaged *G*g in liver weight and *ē*J in oxygen produced per milligram of glycogen in its liver. Therefore, the amount of energy stored in the liver of the large yellow croaker can be determined as follows:

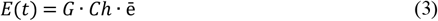

Then, substituting formula (2) into formula (3):

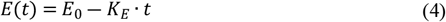

In the formula, *E_0_* is the energy storage of large yellow croaker liver glycogen at the initial moment, *E_0_* = 17.9 *G* · *ē*. The rate of change of liver glycogen storage is *K_E_*=*G* · *Kv*. The average liver weight of large yellow croaker *G* is constant, as is the aerobic metabolism capacity of liver glycogen, so *E_0_* is a constant. Due to *Kv* being related to swimming speed, the change rate of energy storage *K_E_* is also related to swimming speed. It is theorized that when large yellow croaker have exhausted (or nearly exhausted) their liver glycogen storage, their maximum swimming time will be:

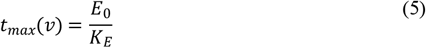

Multiply both sides of equation (4) by *E_0_* to get:

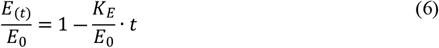

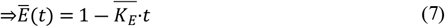

Based on (1), (5), and (7), and based on the previous results, we can calculate the distribution curve of the physical fitness of large yellow croakers based on a liver glycogen consumption model under different conditions of swimming speeds. Figure 7 illustrates the relationship. Formula (1) can be used to calculate the longest continuous period time that a large yellow croaker will swim at different speeds.

**Fig.7.**
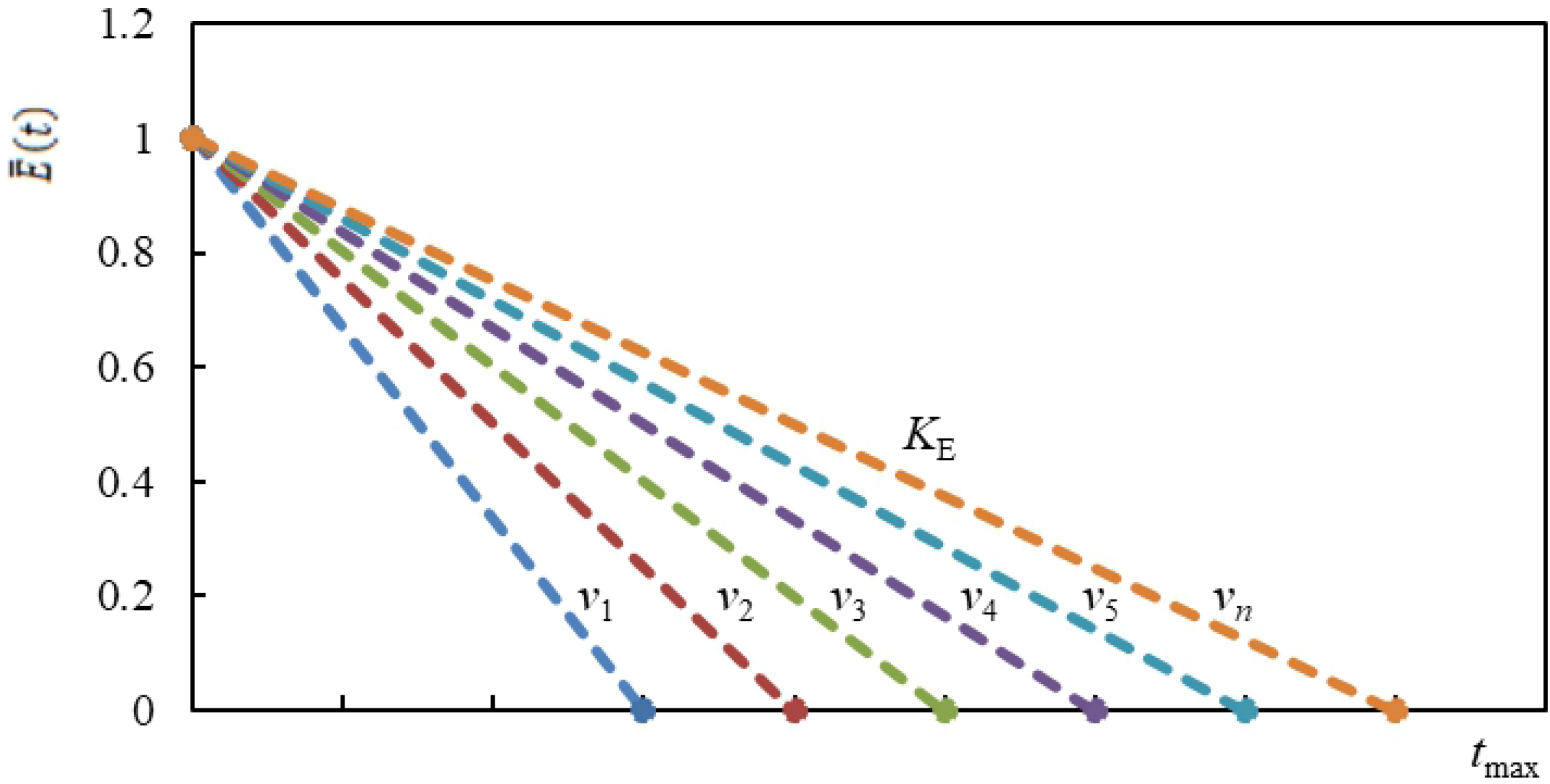
Relationship between swimming ability at different flow velocities

## 4. Discussion

### 4.1 Swimming speed preference of large yellow croakers in clusters

As large yellow croakers have a group activity factor, flow rate preference measurements of single-tailed fish have no direct value for traditional fisheries. Research shows that group swimming affects the metabolic rate of fish and the behavior of the fish while swimming[45]. According to the results, there are significantly more large yellow croakers in the environment with flow velocity below 0.1 m s^-1^ than the other environments. In the experiment, American red carp exhibited similar swimming preferences, but Black breams were scattered at low flow velocity and gathered at 0.42 m s^-1^ flow velocity. This shows that black sea bream grown at a flow rate of 42 cm s^-1^ is optimal for growing[46]. Table 4 summarizes the swimming speed preferences of some fish species.

**Tab.4.** Summary of favorite flow rates of some fish species

| species | body length<br>(cm) | favorite flow rate ($\text{m s}^{-1}$) |
| --- | --- | --- |
| *Megalobrama skolkovii* | 10-17 | 0.3-0.5 |
| *Carassius auratus auratus* | 10-15 | 0.3-0.6 |
| *Cyprinus carpio* | 20-25 | 0.3-0.8 |
| *Hypophthalmichthys molitrix* | 10-15 | 0.3-0.5 |
| *Ctenopharyngodon idella* | 15-18 | 0.3-0.5 |

In this experiment, there are a few fishes distributed at high flow velocity. The experiment process may be affected by environmental factors such as sound or light. There is also the phenomenon of fish catching the net. Based on the result that there is no significant difference in individual specifications (P<0.05), reasons for contacting the internet include a lack of fitness or poor health. This experiment might be used as a screening method to determine the health of fish. Distinguishing the large yellow croaker in inconsistent state in time can avoid the transmission of unknown diseases, and the poor state can also be reared alone to restore the state quickly and reduce losses.

### 4.2 The endurance swimming research of large yellow croaker

At a flow rate of 50 cm s^-1^, the juvenile large yellow croaker’s average endurance time is 20 minutes, according to 5 flow rate experiments. This result provides a reference for the speed of large yellow croaker trawl transportation by fishing vessels. In contrast, Atlantic salmon endurance swimming speed is 0.96-1.02 m s^-1^ [47], the shortnose sturgeon with a body length of 23.8 cm lasts for 170-200 minutes at 0.35 m s^-1^ [48]. the critical swimming speed of American redfish reaches 80 cm s^-1^, the critical swimming speed of bass (*Lateolabrax japonicus*) and oblique mustache (*Hapalogenys nitens*) is 60 cm s^-1^. These fish indicate that large yellow croakers do not have strong swimming abilities.

The large yellow croaker has an elongated body, flat sides, large gaps, the upper jaw is equal to the lower jaw, and its caudal fin is wedge-shaped. In comparison to long fusiform and spindle-shaped fish, their innate body size determines their weak swimming ability. Based on the premise of excluding species differences from the endurance swimming experiment, the differences in body length and weight of the 6 experimental fish groups were measured. The results were not significantly different (P>0.05), exclude Reynolds number influence[49]. The experiment found that when the large yellow croaker swims close to the front of the experimental device when it is full of energy in the initial stage, and constantly adjusts its position in a small range, it shows that it is looking for the most energy-saving spot. As time passed, the large yellow croaker gradually moved to the rear end of the device, struggling to escape when its tail touched the cage, and repeated this many times until it was exhausted and completely leaned against the cage.

### 4.3 The relationship between the swimming ability of large yellow croaker and its metabolites

The swimming process of fish is dominated by aerobic respiration, and the energy source mainly consumes glycogen and blood sugar. It must exercise for a long time at low intensity to consume fat and protein. For this experiment, only blood glucose and glycogen consumption are considered. Despite the continuous replenishment of blood sugar in the fish, the experiment did not directly measure changes in blood sugar content but directly measured glucagon levels. Finally, the glucagon content of the experimental fish increased by about 140 ng L^-1^. In the current research on whether Cori plays a role in fish, some studies report that both lactic acid and hydrogen seem to remain in the white muscle for in situ metabolism. However, the entire metabolic process is still controversial[50, 51]. To calculate the results, the PCA method was used to determine the content of related metabolites. Graphs show that in the large yellow croaker, the changes in the content of lactic acid in the muscle and liver are highly consistent, and the changes in the content of glycogen in the muscle and liver are also highly consistent. The results indicate that the Cori cycle plays a significant role in the body of large yellow croaker. In addition, large yellow croakers do not have strong swimming ability. Except for reasons such as the shape not being suitable for high-speed swimming, the Cori cycle may be longer than the in-situ metabolism process and the energy replenishment speed may be slower. Swimming and exercise abilities differ not only in appearance but also in the entire metabolic mechanism of the body of various species. Compared with the changes of the other two glycogen indicators, there is a much smaller difference between their changes from a numerical perspective. Although adrenaline and glucagon can also break down glycogen, they have a secondary role. The glycogen content in the liver of large yellow croakers decreases with increased swimming time, and the decline is more obvious. It shows that the decrease of blood sugar levels stimulates the secretion of glucagon, which makes liver glycogen break down into blood sugar. As a large yellow croaker becomes fatigued, its liver glycogen reserves are near exhaustion. As a result, when liver glycogen is not able to supply blood sugar stability, but the conversion of fat to sugar will take a long time. In this stage, the blood sugar level drops and hypoglycemia occurs, causing fatigue or loss of exercise capacity. It is speculated that liver glycogen is a key factor affecting its fatigue. There is a fairly mature model of the liver, including glycogenolysis, glycogen production, glycolysis, and lipolysis[52].

The peak value of lactic acid accumulation in fish after exhaustion is an important indicator of the fish’s anaerobic exercise capacity, which explains the significant increase in muscle lactic acid in this experiment. There is a strong correlation between the changing trend in blood lactic acid and the changing trend in muscle lactic acid and a “lactate release” mechanism may be at work in large yellow croaker[53]. This means the Cori cycle exists in large yellow croakers. However, some studies oppose the argument that lactic acid can be exchanged in muscle and blood without obstacles. During its exercise, fish store most of their lactic acid in white muscle, where it is oxidized or undergoes gluconeogenesis[54]. This conclusion is basically consistent with the experimental data of this research. And experimental data in this study support this conclusion. In Figure 5, the ratio of lactic acid in muscle to blood is 1:1000. The study results are almost the same as the change in lactic acid in salmon blood after swimming fatigue. In contrast to benthic species (such as flounder), it releases almost no lactic acid, and the blood lactic acid level is extremely low[55, 56]. The extent of the effects of lactic acid on fish fatigue is still debated, and the experiment did not determine the maximum lactic acid content in the muscles of large yellow croakers. Therefore, muscle lactic acid cannot be regarded as a key factor for fatigue in large yellow croaker juveniles, but it can be used as an influencing factor.

Currently, the energy stored in body fat and the body composition of animals can be measured experimentally. For example, analyzing bioelectrical impedance or isotope water dilution[57]. However, it is impossible to use fats as direct energy sources. Muscle glycogen is a direct energy substance for the movement of large yellow croaker. Its content changes are regulated by complex hormones, and it is difficult to get a good regularity due to the uneven content in the muscles. In this case, an energy distribution model based on yellow croaker fatigue is closer to reality.

Chao Shuai hypothesized that hepatic glycogen was used as an energy source to build a physical fitness model for American red fish for endurance swimming. However, liver glycogen has much less energy than fat and muscle, and the explanation for it as an energy supply substance is not sufficient. Based on the results of the chemical indicators, it appears that liver glycogen consumption is indeed related to fatigue in juvenile large yellow croakers, so a model based on liver glycogen content makes sense. This article constructs a model from the perspective of changes in blood glucose content since liver glycogen is mainly used to regulate blood glucose concentration. The model is based on the assumption that the liver glycogen of juvenile large yellow croaker serves as the only source for blood sugar supplementation during continuous swimming; at the same time, the individual differences of large yellow croakers are considered not significant (P<0.05).

Large yellow croaker is mainly farmed in cages, deep-water cages, however, these farming methods are limited by several factors such as technical difficulty, cost, and the inability to withstand wind or waves. It has good body and meat quality, color, and body shape when cultured in deep water cages. The results of this experiment showed that large yellow croaker groups prefer environments where the water velocity is less than 10 cm/s, and their endurance swimming ability is expressed using the logarithm: *F*(*x*) = 754.98 − 1508,55*x*. To maintain blood sugar balance, construct a swimming ability model that includes liver glycogen: 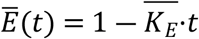. This model considered only the influence of flow velocity as a single parameter. In relevant studies, it has been shown that the estimation of swimming speed and fatigue time by using a closed swimming room could underestimate the actual swimming ability of fish[58, 59], and a practical application is still a long way off. As a potential fatigue factor, it is presently only being studied in the laboratory. The next stage will be to sample the actual sea area to improve the model data and provide detailed information on the selection of deep-sea cage breeding sites for large yellow croakers. The following findings and suggestions are based on this experiment:

1. The flow rate can be set in the breeding pond for large yellow croakers farmed in factories in order to prevent them from hitting the wall and being injured or killed.
2. It is usually designed according to the durability of the fish’s swimming speed[60], and certain requirements are put forward for the continuity and velocity of the water flow. The fishway can be designed with different flow rates, and the fastest flow rate can be adjusted according to the preferences of the fish.
3. Fish fatigue is related to anaerobic respiration in the body of the fish that occurs when it is fatigued and continues to swim for 200 minutes in a row. Lactic acid builds up in the body of the fish, making it feel exhausted. The contents of glycogen were determined in this experiment, and the results showed that when the liver glycogen was exhausted, it was close to the time when the juvenile large yellow croaker reached the net, indicating that it was not able to maintain blood sugar levels in a timely manner. It is speculated that blood glucose stability provides a key role in maintaining the physical fitness of large yellow croaker juveniles.
4. This study provides a new method for constructing the endurance swimming ability model of large yellow croakers and provides a reference for the construction of endurance swimming models for other species of fish.

## Notes

### Competing Interest Statement

The authors have declared that no competing interests exist.

